# Structure of V-ATPase from citrus fruit

**DOI:** 10.1101/2022.05.01.490144

**Authors:** Yong Zi Tan, Kristine A. Keon, Rana Abdelaziz, Peter Imming, Waltraud Schulze, Karin Schumacher, John L Rubinstein

**Affiliations:** Molecular Medicine Program, The Hospital for Sick Children Research Institute, Toronto, ON M5G 0A4, Canada; Department of Biological Sciences, National University of Singapore, 16 Science Drive 4, Singapore 117558, Singapore, and Disease Intervention Technology Laboratory, Immunos, Agency for Science, Technology and Research (A*STAR), 8A Biomedical Grove, Singapore 138648, Singapore; Institut Für Pharmazie, Martin-Luther-Universität Halle-Wittenberg, 06120 Halle (Saale), Germany; Department of Plant Systems Biology, University of Hohenheim, 70593 Stuttgart, Germany; Centre for Organismal Studies, Heidelberg University, 69120 Heidelberg, Germany; Department of Medical Biophysics, University of Toronto, Toronto, ON M5G 1L7, Canada; Department of Biochemstry, University of Toronto, Toronto, ON M5S 1A8, Canada

## Abstract

Vacuolar-type ATPases (V-ATPases) are rotary proton pumps involved in numerous essential cellular processes in all eukaryotes. Difficulty in obtaining preparations of purified V-ATPase from plants in sufficient quantities for structural analysis has hindered determining the 3D structure of the plant V-ATPase. We used the *Legionella pneumophila* effector SidK to affinity-purify the endogenous V-ATPase from lemon fruit. The preparation was sufficient to partially cover an electron microscopy specimen grid, allowing structure determination for the enzyme in two rotational states. The structure defines the ATP:H^+^ ratio of the enzyme, demonstrating that it can establish a maximum ΔpH of ∼3, which is insufficient to maintain the low pH observed in the vacuoles of juice sac cells in lemons and other citrus fruit. Compared to the yeast and mammalian enzymes, the membrane region of the plant V-ATPase lacks subunit f and possesses an unusual configuration of transmembrane α helices. Subunit H, which inhibits ATP hydrolysis in the isolated catalytic region of V-ATPase, adopts two different conformations in the intact complex, hinting at a role in modulating activity in the intact enzyme.

## Introduction

V-ATPases are rotary enzyme complexes that couple the energy released from hydrolysis of adenosine triphosphate (ATP) to the translocation of protons across a membrane. Ubiquitous in eukaryotes, V-ATPase activity acidifies the lumen of various organelles, enabling pH-dependent processes and driving transmembrane transport (Forgac, 2007; Kane, 2007; Vasanthakumar and Rubinstein, 2020). Understanding of the structure-function relationship in V-ATPases has been driven by studies of the enzyme from yeast (Benlekbir et al., 2012; Mazhab-Jafari et al., 2016; Oot et al., 2016; Roh et al., 2018; Vasanthakumar et al., 2019, 2022; Zhao et al., 2015), mammals (Abbas et al., 2020; Tan et al., 2021; Wang et al., 2020a, 2020b), and insects (Muench et al., 2009). These studies revealed the overall architecture of the enzyme and the sequence of events that couples ATP hydrolysis to proton pumping. ATP hydrolysis in the hexameric A_3_B_3_ subcomplex of the soluble catalytic V_1_ region causes the three pairs of A and B subunits to cycle between “Open”, “Loose”, and “Tight” conformations (Forgac, 2007; Kane, 2007; Zhao et al., 2015). These conformational changes induce rotation of the rotor subcomplex, consisting of subunits D, F, d, and a ring of c subunits in the membrane-embedded V_O_ region of the complex. In yeast, the c-ring contains eight c subunits, a c′ subunit, and a c″ subunit (Mazhab-Jafari et al., 2016) while in mammals it contains nine c subunits and a c″ subunit (Abbas et al., 2020). In yeast, the c-ring surrounds transmembrane α helices from subunit c″ (Mazhab-Jafari et al., 2016) and Voa1p (Roh et al., 2018), while in mammals it surrounds transmembrane α helices from subunit c″, subunit ATP6AP1/Ac45, and subunit ATP6AP2/PRR (Abbas et al., 2020). Rotation of the c-ring against the membrane-embedded C-terminal domain of subunit a in V_O_ drives proton translocation across the membrane. Subunit a also interacts with the membrane-embedded subunits e and f (Mazhab-Jafari et al., 2016), neither of which have a known function beyond helping to anchor the static part of the complex in the membrane. Subunit f is also known as RNAseK in mammals (Abbas et al., 2020). The soluble N-terminal domain of subunit a connects to a collar region consisting of subunits H and C, as well as three peripheral stalks, each consisting of a heterodimer of subunits E and G. These three peripheral stalks prevent subunit a from rotating with the rotor during ATP hydrolysis in V_1_.

V-ATPase activity in yeast (Kane, 1995), insects (Sumner et al., 1995), and mammals (Liberman et al., 2014; Sautin et al., 2005; Stransky and Forgac, 2015) is regulated by reversible separation of the V_1_ region from the V_O_ region. On separation, the V_O_ complex adopts an inhibited conformation (Mazhab-Jafari et al., 2016) that is impermeable to protons (Ward and Sze, 1992; Zhang et al., 1992) and the V_1_ complex undergoes a major conformational rearrangement that blocks ATP hydrolysis and prevents its premature reassembly with V_O_ (Vasanthakumar et al., 2022). Inhibition of ATP hydrolysis in dissociated V_1_ requires subunit H (Parra et al., 2000), which forms a new interaction with one of the peripheral stalks on dissociation to prevent the conformational changes needed for hydrolysis (Vasanthakumar et al., 2022). Subunit C subsequently separates from V_1_ (Gräf et al., 1996), with the RAVE complex (regulator of the <u>A</u>TPase of <u>v</u>acuoles and <u>e</u>ndosomes) required for reassembly of V_1_, V_O_, and subunit C to form a functional V-ATPase (Jaskolka et al., 2021; Smardon et al., 2002).

In plants, V-ATPases are found at the membrane limiting the vacuole or tonoplast, and in the trans-Golgi network/early endosome of all cells (Schumacher, 2014). The presence of a large central vacuole that occupies up to 90% of the cell volume is a hallmark of plant cells that allows plants to increase the surface of their photosynthesizing and nutrient-absorbing organs with minimal cost. Beside filling space, vacuoles act as the main storage compartment for solutes and essential nutrients, help to detoxify the cytosol, and serve as lysosome-like organelles in which endocytic and autophagic cargo are digested (Martinoia et al., 2012; Marty, 1999). All vacuolar functions require massive fluxes of molecules across the tonoplast, which are energized by the proton gradient and membrane potential. It has long been assumed that the combined activity of V-ATPase and the vacuolar H+-pyrophosphatase (V-PPase) enables plants to maintain transport into the vacuole even under stressful conditions (Maeshima, 2000). However, whereas vacuolar pH is 5 to 6 in typical cells, certain plant cell types, such as lemon fruit juice sac cells have much more acidic vacuoles with pH ∼2 (Echeverria and Burns, 1989). V-ATPases in yeasts and mammals pump protons with an ATP:H^+^ ratio of 3:10 and thus can only produce a ΔpH of ∼3 (Abbas et al., 2020; Mazhab-Jafari et al., 2016), which would be incapable of producing the vacuolar pH of ∼2 seen in citrus fruits. However, citrus belongs to a group of phylogenetically unrelated plant species that contain an additional vacuolar proton-pumping P-ATPase complex consisting of PH5, a P3A-ATPase, and PH1, a P3B-ATPase similar to bacterial Mg transporters (Li et al., 2016), and the sour taste of citrus fruits correlates with the expression levels of these proteins (Strazzer et al., 2019).

Negative stain electron microscopy, which can provide low-resolution protein structures from extremely small quantities of purified material, showed the plant V-ATPase from *Kalanchoë daigremontiana* resembles other V-ATPases (Domgall et al., 2002). To determine if plant, and in particular citrus, V-ATPases possess features that differ from other V-ATPases, we isolated the enzyme from lemon fruits and subjected the protein to analysis by electron cryomicroscopy (cryoEM). The structure reveals that the citrus V-ATPase maintains the same ATP:H^+^ ratio as other V-ATPases, but lacks subunit f and has an unusual arrangement of transmembrane α helices in the middle of its c-ring. Further, the inhibitory subunit H from the V_1_ region can adopt a conformation not seen previously in intact V-ATPase structures, which could suggest an additional role in enzyme regulation.

## Results and Discussion

### Overall structure of the plant V-ATPase

Previous structural studies of yeast V-ATPase have relied on purification of the detergent-solubilized endogenous enzyme following incorporation of sequence encoding an affinity tag into the yeast chromosomal DNA (Benlekbir et al., 2012; Mazhab-Jafari et al., 2016; Oot et al., 2016; Roh et al., 2018; Vasanthakumar et al., 2022; Zhao et al., 2015). V-ATPase from bovine brain, where the enzyme is highly enriched, was isolated for structural studies using a large mass of starting material and conventional chromatographic methods (Wang et al., 2020b). However, the high-affinity interaction between the *Legionella pneumophila* effector SidK and V-ATPase (Sharma and Wilkens, 2017; Xu et al., 2010; Zhao et al., 2017) has also allowed affinity purification of the complex from smaller amounts of mammalian tissue (Abbas et al., 2020). With this approach, a SidK fragment comprising residues 1-278 of the protein is expressed recombinantly bearing a C-terminal 3×FLAG tag, immobilized on an anti-FLAG affinity column, and used to bind V-ATPase from detergent-solubilized membranes. The SidK_1-278_:V-ATPase complex is then eluted from the column with 3×FLAG peptide. Despite SidK usually binding mammalian V-ATPase during infection by *L. pneumophila*, the interaction is preserved even with V-ATPase from the evolutionarily distant *Saccharomyces cerevisiae* (Maxson et al., 2022; Zhao et al., 2017). Therefore, for structural analysis of the V-ATPase from lemon fruits we attempted purification using SidK.

Initially, we isolated V-ATPase from eight lemons. While this preparation resulted in sufficient enzyme for negative stain electron microscopy (EM) (Fig. 1A-i), the concentration was insufficient for electron cryomicroscopy (cryoEM), which can require sample that is 100-fold more concentrated. We subsequently purified V-ATPase from eighty fruits and concentrated it with a 100 kDa molecular weight cut-off centrifuged contractor device. This preparation yielded sufficient sample to perform mass spectrometry and partially cover two cryoEM grids with sample. It was insufficient for performing SDS-PAGE or an activity assay. In the electron microscope, most of the specimen grid was observed to be dry due to the small volume of sample applied. CryoEM images from regions that had ice (Fig. 1A-ii) showed a low density of protein particles, owing to the relatively low protein concentration of the sample. Nonetheless, automated data acquisition allowed collection of 12,974 micrographs, which provided 180,946 particle images. Two-dimensional (2D) classification of these images produced class averages that reveal the characteristic V_1_ and V_O_ regions of V-ATPase (Fig. 1A-iii).

**Figure 1.**
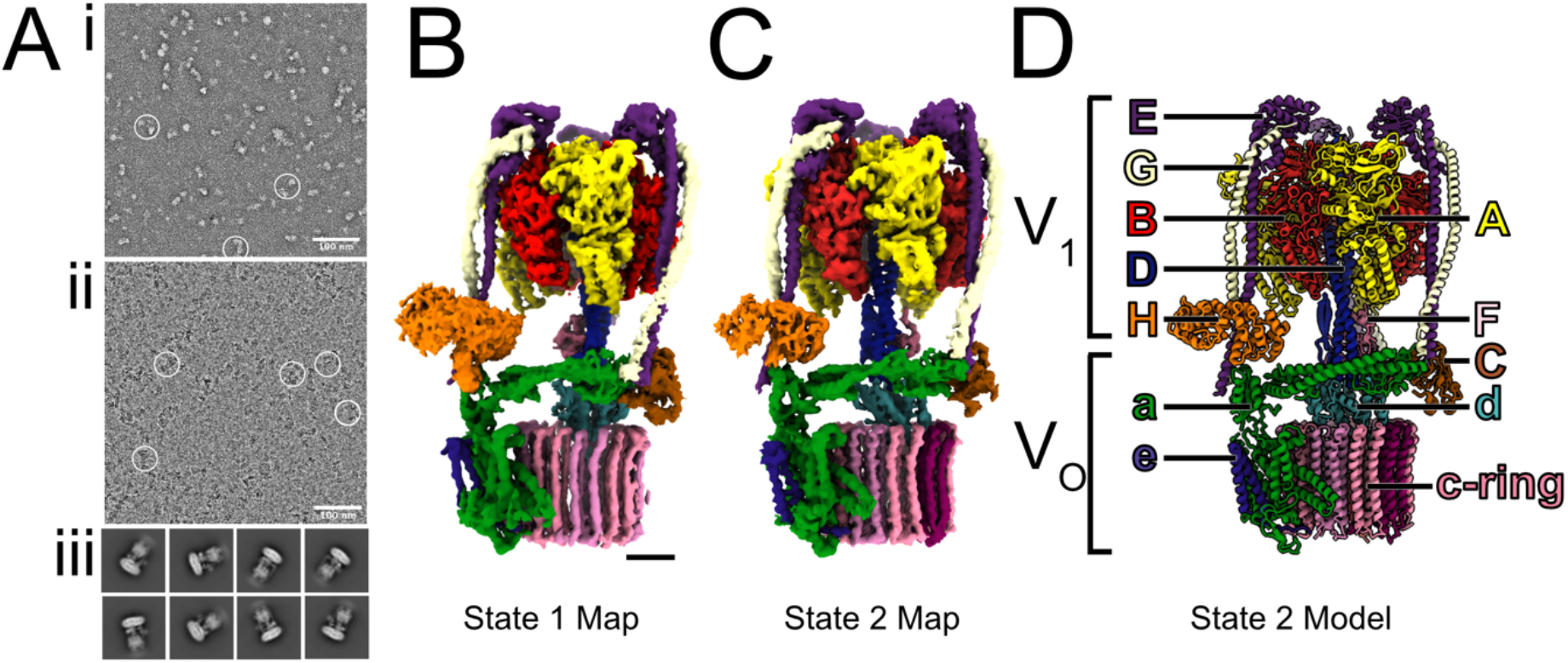
Electron microscopy of the citrus V-ATPase. **A**, Representative negative stain electron micrograph (i), cryoEM micrograph (ii), and 2D class average images (iii) from cryoEM of citrus V-ATPase. Example particles are circled in white in the micrographs. **B**, CryoEM map of the citrus V-ATPase in rotational State 1. Scale bar, 25 Å. **C**, CryoEM map of the citrus V-ATPase in rotational State 2. **D**, Atomic model of the citrus V-ATPase in rotational State 2.

Typically, when imaged with cryoEM, intact V-ATPase adopts three rotary states related by a ∼120° rotation of the rotor and known as rotational “State 1”, “State 2”, and “State 3” (Zhao et al., 2015). Three-dimensional (3D) reconstruction and classification with the dataset of images of citrus V-ATPase led to structures of the enzyme in rotary States 1 and 2, with nominal overall resolutions of 4.0 Å and 3.9 Å, respectively (Fig. 1B and C, Supplementary Fig. 1). Subsequent focused refinement of the V_1_ and V_O_ regions of the maps improved the resolution for these regions of the complex (Supplementary Fig. 1C). Surprisingly, the SidK used to purify V-ATPase was absent in the maps, suggesting that the *Legionella* effector has a lower affinity for the plant V-ATPase than for mammalian V-ATPase and can dissociate during purification or specimen preparation.

To identify subunits of the citrus V-ATPase in the sample, the detergent-solubilized protein was digested in solution with trypsin and liquid-chromatography tandem mass spectrometry (LC-MS/MS) was performed. Spectra were matched against the proteomes of *Citrus clementina* and *Citrus sinensis*, and also sequenced de-novo (see methods). V-ATPase subunits were then identified by comparison to the well-annotated *Arabidopsis thaliana* proteome (Supplementary Data 1). This analysis detected V-ATPase subunits A, D, E1, G, H, and a3. The sequences for these subunits were used to generate initial models of the subunits with RoseTTAfold (Baek et al., 2021) and PHYRE2 (Kelley et al., 2015), which were compared and the better-fitting model placed in cryoEM map and refined. In addition, the citrus orthologs of *Arabidopsis* subunits B2, C, F, c″, d2, e1, AP1, and AP2 were modelled, fit, and refined into the maps (Fig. 1D, Table 1). In general, the resolution of the maps was insufficient for reliable modelling of amino acid side chain orientations and side chains were truncated to Cα, except within regions in the A_3_B_3_ subcomplex and some bulky residues in subunits E, G, and e (Supplementary Fig. 2).

**Table 1:**
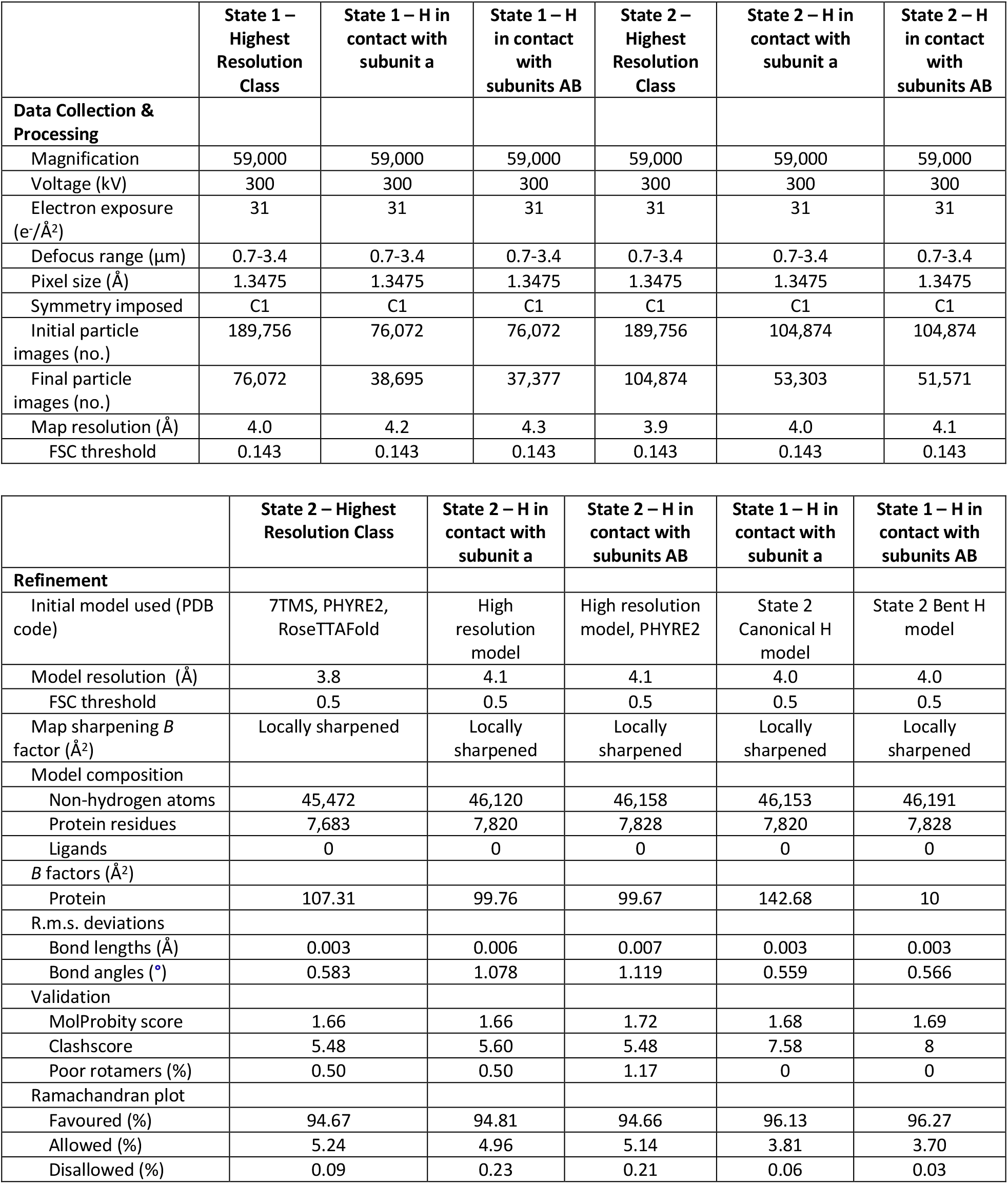
CryoEM map validation and model building statistics.

|  | State 1 – Highest Resolution Class | State 1 – H in contact with subunit a | State 1 – H in contact with subunits AB | State 2 – Highest Resolution Class | State 2 – H in contact with subunit a | State 2 – H in contact with subunits AB |
| --- | --- | --- | --- | --- | --- | --- |
| **Data Collection & Processing** |  |  |  |  |  |  |
| Magnification | 59,000 | 59,000 | 59,000 | 59,000 | 59,000 | 59,000 |
| Voltage (kV) | 300 | 300 | 300 | 300 | 300 | 300 |
| Electron exposure (e<sup>-</sup>/Å<sup>2</sup>) | 31 | 31 | 31 | 31 | 31 | 31 |
| Defocus range (μm) | 0.7-3.4 | 0.7-3.4 | 0.7-3.4 | 0.7-3.4 | 0.7-3.4 | 0.7-3.4 |
| Pixel size (Å) | 1.3475 | 1.3475 | 1.3475 | 1.3475 | 1.3475 | 1.3475 |
| Symmetry imposed | C1 | C1 | C1 | C1 | C1 | C1 |
| Initial particle images (no.) | 189,756 | 76,072 | 76,072 | 189,756 | 104,874 | 104,874 |
| Final particle images (no.) | 76,072 | 38,695 | 37,377 | 104,874 | 53,303 | 51,571 |
| Map resolution (Å) | 4.0 | 4.2 | 4.3 | 3.9 | 4.0 | 4.1 |
| FSC threshold | 0.143 | 0.143 | 0.143 | 0.143 | 0.143 | 0.143 |

|  | State 2 – Highest Resolution Class | State 2 – H in contact with subunit a | State 2 – H in contact with subunits AB | State 1 – H in contact with subunit a | State 1 – H in contact with subunits AB |
| --- | --- | --- | --- | --- | --- |
| **Refinement** |  |  |  |  |  |
| Initial model used (PDB code) | 7TMS, PHYRE2, RoseTTAFold | High resolution model | High resolution model, PHYRE2 | State 2 Canonical H model | State 2 Bent H model |
| Model resolution (Å) | 3.8 | 4.1 | 4.1 | 4.0 | 4.0 |
| FSC threshold | 0.5 | 0.5 | 0.5 | 0.5 | 0.5 |
| Map sharpening *B* factor (Å<sup>2</sup>) | Locally sharpened | Locally sharpened | Locally sharpened | Locally sharpened | Locally sharpened |
| Model composition |  |  |  |  |  |
| Non-hydrogen atoms | 45,472 | 46,120 | 46,158 | 46,153 | 46,191 |
| Protein residues | 7,683 | 7,820 | 7,828 | 7,820 | 7,828 |
| Ligands | 0 | 0 | 0 | 0 | 0 |
| *B* factors (Å<sup>2</sup>) |  |  |  |  |  |
| Protein | 107.31 | 99.76 | 99.67 | 142.68 | 10 |
| R.m.s. deviations |  |  |  |  |  |
| Bond lengths (Å) | 0.003 | 0.006 | 0.007 | 0.003 | 0.003 |
| Bond angles (°) | 0.583 | 1.078 | 1.119 | 0.559 | 0.566 |
| Validation |  |  |  |  |  |
| MolProbity score | 1.66 | 1.66 | 1.72 | 1.68 | 1.69 |
| Clashscore | 5.48 | 5.60 | 5.48 | 7.58 | 8 |
| Poor rotamers (%) | 0.50 | 0.50 | 1.17 | 0 | 0 |
| Ramachandran plot |  |  |  |  |  |
| Favoured (%) | 94.67 | 94.81 | 94.66 | 96.13 | 96.27 |
| Allowed (%) | 5.24 | 4.96 | 5.14 | 3.81 | 3.70 |
| Disallowed (%) | 0.09 | 0.23 | 0.21 | 0.06 | 0.03 |

### Citrus V-ATPase structure sets the ATP:H^+^ ratio at 3:10

The overall structure and subunit composition of the citrus V-ATPase resembles the yeast and mammalian enzymes (Abbas et al., 2020; Mazhab-Jafari et al., 2016; Roh et al., 2018; Zhao et al., 2015). The V_1_ region contains subunits A_3_B_3_CDE_3_FG_3_H (Fig. 1D). However, the V_O_ region has notable differences with the yeast and mammalian enzymes (Fig. 2). The citrus V_O_ region comprises subunits acc″de, AP1, and AP2 (Fig. 2A-i). The citrus V_O_ lacks subunit f, a subunit of unknown function that was identified in yeast V-ATPase as the protein YPR170W-B (Mazhab-Jafari et al., 2016) (Fig. 2A-ii) and RNAseK in mammals (Fig. 2A-iii) (Abbas et al., 2020). Additionally, in contrast to the yeast enzyme, which contains subunit c′ (Fig. 2A-ii), the c-ring in citrus is composed of nine c subunits and one c″ prime subunit (Fig. 2A-i), as in mammals (Fig. 2A-iii).

**Figure 2.**
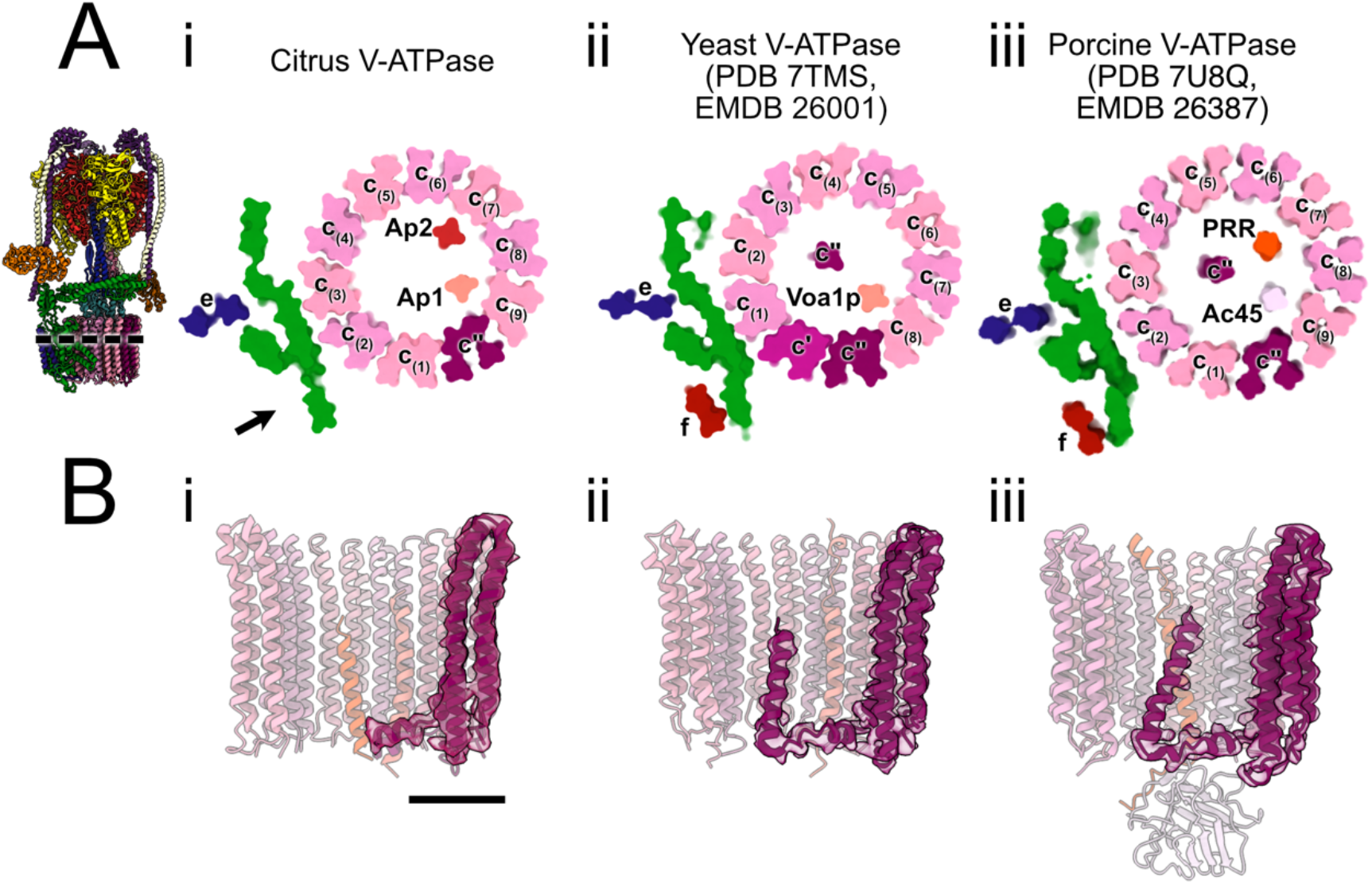
Comparison of the citrus, yeast, and porcine V-ATPase. **A**, Transverse cross sections through the V_O_ region from citrus (i), yeast (ii), and porcine (iii) V-ATPases. Blue arrow indicates the position occupied by subunit f in other V-ATPases. **B**, Longitudinal cross sections through the c-ring showing subunit c″ in the citrus (i), yeast (ii), and porcine (iii) V-ATPases, rendered in magenta. Scale bar, 25 Å.

The c and c″ subunits in V_O_ are each capable of transporting a single proton across the membrane during rotation of the ring past subunit a during catalysis. A 360° rotation of the rotor results in translocation of 10 protons and hydrolysis of three ATP molecules. Therefore, the c-ring stoichiometry of the citrus enzyme sets the ATP:H^+^ ratio for the complex and allows determination of the maximum ΔpH that can be established by the complex from the free energy of hydrolysis of ATP (Nicholls and Ferguson, 2013). 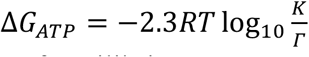, where *R* is the gas constant, *T* is the temperature, and *K*/*Γ* is the ratio of equilibrium constant to mass action ratio for ATP hydrolysis in the cytoplasm. At a typical cytoplasmic *K*/*Γ* value of 10^10^ (Nicholls and Ferguson, 2013) and a temperature of 25°C, this equation gives Δ*G*_*ATP*_ = −57 *kJ*/*mol*. The V-ATPase converts this free energy into a free energy of proton translocation, 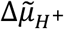. With an ATP: H^+^ ratio of 3:10, the V-ATPase can use the free energy of ATP hydrolysis to establish 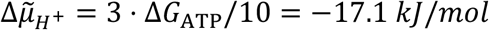. This 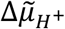 can be composed of Δ*ψ*, the electrical potential across the membrane, and ΔpH, the pH difference across the membrane, according to the relationship:

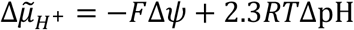

where *F* is the Faraday constant. If the V-ATPase is the only electrogenic pump at the membrane, the ΔpH is maximized at Δ*ψ* = 0, which gives a ΔpH = 3. Therefore, V-ATPase alone cannot account for the extreme ΔpH seen in the citrus vacuole. Instead, this role is likely filled by a P-type ATPase, with the P-type ATPases PH1 and PH5 having been identified previously as contributing to maintenance of the acidified vacuole of citrus fruits (Strazzer et al., 2019). Following the analysis from above, the ATP:H^+^ ratio of 1 in P-type ATPases allows a ΔpH of up to 10, enabling increased acidification of the vacuole relative to the cytoplasm at a much higher bioenergetic expense than V-ATPase. Therefore, it appears that V-ATPase can reduce the vacuole pH by 3 units relative to the cytoplasm, possibly by recruitment of V-ATPases to the increasingly acidic pre-vacuolar compartments (Maxson et al., 2022), with the P-type ATPases achieving the ultimate pH of the vacuole.

### Plant V-ATPase possesses a c-ring that differs from the mammalian and yeast rings

In addition to the 40 transmembrane α helices that make up the ring of c subunits, two additional transmembrane α helices are enclosed within the middle of the citrus c-ring (Fig. 2A-i). The yeast c-ring also encloses two additional transmembrane α helices (Fig. 2A-ii) while the mammalian c-ring encloses three (Fig. 2A-iii). However, in both the yeast and mammalian enzyme, one of these additional transmembrane α helices is an N-terminal α helix from subunit c″ (Fig. 2B, *center* and *right*). Strikingly, the citrus c″ subunit (182 residues), which is shorter than the yeast and mammalian c″ subunits (e.g. 213 residues in *S. cerevisiae* and 205 residues in *Sus scrofa*) lacks sufficient sequence to form this transmembrane α helix (Fig. 2B, *left*). The first transmembrane α helix in the middle of the citrus c-ring appears to be from the protein AP1 (Fig. 2A-i, *peach*), a homolog of the yeast protein Voa1p (Fig. 2A-ii, *peach*) and a portion mammalian ATP6AP1/Ac45 (Fig. 2A-iii, *orange-red*), which also form transmembrane α helices in those enzymes. The second transmembrane α helix in the middle of the c-ring corresponds to AP2, a homologue of ATP6AP2/PRR in mammalian V-ATPase (Fig. 2A-iii, *orange-red*) a fragment of the (pro)renin receptor, which is absent in *S. cerevisiae* (Fig. 2-Aii).

While insect and mammalian V-ATPases possess a large domain of unknown function from subunit ATP6AP1/Ac45 that extends into the lumen of the acidified compartment, the yeast V-ATPase homologue Voa1p is processed into a single short transmembrane span (Roh et al., 2018). Voa1p in yeasts, and likely ATP6AP1/Ac45 in mammals, is involved in assembly of the V_O_ region of V-ATPase (Ryan et al., 2008). Plant V-ATPases appear to require a protein like Voa1p for the same function, as seen by the placement of AP1 in the structure in a position equivalent to Voa1p and ATP6AP1/Ac45 in yeast and mammals, respectively. The citrus AP1 resembles yeast Voa1p (Roh et al., 2018), lacking the large luminal domain of unknown function seen in ATP6AP1/Ac45 (Abbas et al., 2020) (Fig. 2B). Similarly, AP2 in the citrus enzyme appears in the same position as ATP6AP2/PRR in the mammalian enzyme (Fig. 2A). However, the absence of a yeast homologue for AP2 or ATP6AP2/PRR suggests that the presence of this protein within the c-ring is a feature only in V-ATPases from higher eukaryotes such as mammals and plants.

### Subunit H may inhibit intact plant V-ATPase

Subunit H in the V_1_ region of V-ATPase is responsible for inhibition of ATPase activity when V_1_ separates from V_O_ (Parra et al., 2000). The N-terminal domain of subunit H is bound to peripheral stalk #1 of the V-ATPase (Fig. 3A, “*PS #1*”). On dissociation of V_1_ from V_O_, the C-terminal domain of subunit H loses contact with subunit a in V_O_ and binds the second peripheral stalk of the enzyme, preventing conformational changes in V_1_ and inhibiting ATP hydrolysis (Vasanthakumar et al., 2022). Density for the C-terminal domain of subunit H in the 3D maps of citrus V-ATPase was missing for both States 1 and 2 (Fig. 3A, *circled in blue*). However, 3D variability analysis of particle images (Punjani and Fleet, 2021) from both rotary states revealed that the C-terminal domain of subunit H could adopt two different conformations, with the consensus map missing the domain due to incoherent averaging. In one extreme of the principal component analysis, subunit H adopts the conformation seen in previous structures of intact V-ATPase (Benlekbir et al., 2012; Tan et al., 2021; Vasanthakumar et al., 2019, 2022; Wang et al., 2020a, 2020b; Zhao et al., 2015), with the C-terminal domain of H bound to the N-terminal domain of subunit a (Fig. 3B and C). In the other extreme, the C-terminal domain undergoes a rotation of 130° in State 2 and 110° in State 1 to bind an AB pair from the V_1_ region, inserting between the two subunits (Fig. 3B and D, Video 1). In rotational State 2 (Fig. 3D-iii) the AB pair adopts a more open conformation than in rotational State 1 (Fig. 3D-iv) and the C-terminal domain of H binds deeper in the cleft between the two subunits to maintain its interaction (Video 2).

**Figure 3.**
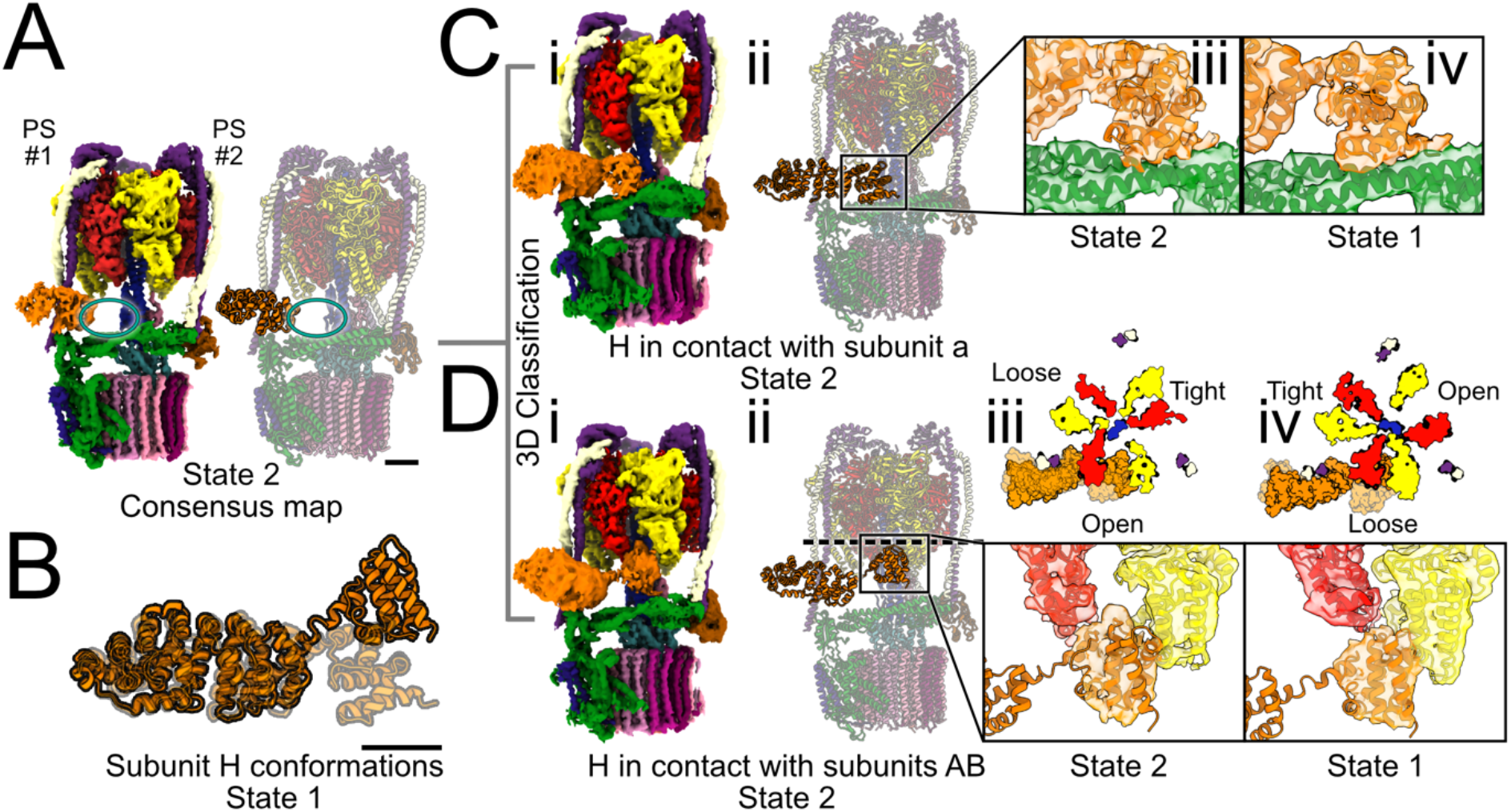
Multiple conformations of subunit H in the citrus V-ATPase. **A**, State 2 consensus map with C-terminal domain of subunit H missing (*blue circle*). PS #1, peripheral stalk #1. PS #2, peripheral stalk #2. **B**, Overlay of two subunit H conformations from rotational State 1. The conformation that contacts subunit a is shown in faded orange while the conformation that contacts the subunit AB pair is shown in opaque orange. Scale bar, 25 Å. **C**, Map (i) and model (ii) showing subunit H in contact with subunit a for state 2, with closeup views of the interaction between subunit H and subunit a in State 2 (iii) and State 1 (iv). **D**, Map (i) and model (ii) showing H in contact with subunits AB for state 2, with cross-section and closeup views of the interaction shown for State 2 (iii) and State 1 (iv).

The conformation of subunit H that contacts an AB pair resembles a conformation seen in a crystal structure of a V_1_ΔC complex (Oot et al., 2016) where it was proposed to inhibit ATP hydrolysis by V_1_. It was observed again later by cryoEM of a complex that formed from purified V_1_ΔC over several days at 4°C that included V_1_ΔC with two additional H subunits and an additional EG peripheral stalk (Vasanthakumar et al., 2022). In intact citrus V-ATPase, the rotation of the C-terminal domain from the conformation in contact with subunit a to the conformation that contacts an AB pair is less than the 150° rotation observed in the crystal structure. Further, in the structure observed here the C-terminal domain of H does not contact the central rotor subunits D and F. However, the subunit H conformation seen in the V_1_ΔC crystal structure and V_1_ΔC+H_2_EG cryoEM structure suggests that the C-terminal domain of subunit H possesses affinity for the AB interface. It is tempting to speculate that in addition to inhibiting ATP hydrolysis in the dissociated V_1_ complex, subunit H can inhibit activity in intact V-ATPase by bending to contact an AB pair. As described above, the ΔpH of the juice sac cell vacuole is too acidic to have been produced by V-ATPase alone. With 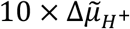 being more negative than 3 × Δ*G*_*ATP*_, the enzyme would be expected to run in reverse and function as an ATP synthase (Guo and Rubinstein, 2022), much as ATP synthases can function as proton pumps when 3 × Δ*G*_*ATP*_ is more negative than 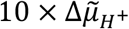. Thus, the unique conformation of subunit H may serve to inhibit ATP synthesis by the citrus V-ATPase when the vacuole is fully acidified, which, if it were to occur, would lead to dissipation of the extreme pH of the juice sac cell vacuole.

Inhibition of the enzyme due to subunit H binding an AB pair is also consistent with rotational State 3 not being detected in the cryoEM dataset. In rotational State 3, the AB pair closest to subunit H is in the “Tight” conformation, which could prevent the C-terminal domain of H from bind between the A and B subunits and lead to the enzyme preferentially adopting States 1 and 2.

Overall, we have demonstrated that affinity purification of V-ATPase from citrus fruit with SidK_1-278_, although providing an extremely limited yield of enzyme, can be used for structure determination by cryoEM. The resulting structure shows that plant V-ATPases have many features in common with yeast and mammalian V-ATPases, but lack subunit f and display marked differences in the enzyme’s c-ring. The enzyme’s ATP:H^+^ ratio puts an upper bound on the ΔpH than can be established by V-ATPase, which is substantially lower than the ΔpH observed across the vacuolar membrane in citrus juice sac cells. Further, we resolved an unusual conformation of subunit H in the citrus V-ATPase. We speculate that this conformation may provide a regulatory mechanism to prevent ATP synthesis by V-ATPase to preserve the extremely low pH of the citrus juice sac cell vacuole. Building on these results, future studies of V-ATPase from the model organism *Arabidopsis thaliana* will reveal if features of citrus V-ATPase described here are unique to citrus or are general attributes of plant V-ATPases.

## Methods

### Purification of V-ATPase from lemon

All steps were performed at 4°C unless otherwise noted. Lemons were purchased from a supermarket and processed eight at a time. The fruit was peeled and the skin discarded. Each peeled fruit was quartered and all eight fruits were blended in a waring blender for 2 min with 430 mL of homogenization buffer (1.5 M 3-(*N*-morpholino)propanesulfonic acid-KOH [MOPS-KOH], pH 8.5, 0.75% bovine serum albumin, 7.5 mM ethylenediaminetetraacetic acid [EDTA], and 0.1 mM phenylmethanesulfonyl fluoride [PMSF]) (Müller et al., 1997). The homogenate was filtered through a stainless-steel kitchen strainer to remove the pulp and large debris. The filtrate was then centrifuged at 800 ×g for 30 min to pellet more debris. The supernatant was then subjected to ultracentrifugation at 200,000 ×g for 30 min to collect cell membranes, yielding an ∼3 g of pellet that was flash frozen and stored at -80 °C. For solubilization of membranes, the pellet was resuspended in solubilization buffer (50 mM 4-(2-hydroxyethyl)-1-piperazineethanesulfonic acid [HEPES] pH 7, 320 mM sucrose, 300 mM NaCl, 10 % [v/v] glycerol, 5 mM EDTA, 5 mM aminocaproic acid, 5 mM para aminobenzamidine, 0.2 mM PMSF) at 10 mL g^-1^ and resuspended with a Dounce homogenizer. Dodecyl maltoside (DDM) was added to 1% (w/v) and the solution was left rocking for 2 h. Insoluble material was removed by centrifugation at 200,000 ×g for 60 min and the supernatant was filtered through a 0.22 μm filter.

Recombinant SidK_1-278_ encoded by plasmid pSAB35 (Addgene plasmid #175787) was expressed in *E. coli* BL21 Codon+ cells and purified as described previously (Abbas et al., 2020) and bound to a column (2 mL) of M2 affinity gel matrix (Sigma). Solubilized lemon membranes were passed through the column to allow V-ATPase to bind. The column was then washed with five column volumes of wash buffer (50 mM HEPES pH 7, 300mM NaCl, 0.025% glycol diosgenin [GDN]) and V-ATPase was eluted with three column volumes of elution buffer (wash buffer with 150 μg mL^-1^ 3×FLAG) and an additional five column volumes of wash buffer. The eluant was concentrated and passed through a Superose 6 Increase 10/300 gel filtration column (GE Healthcare). The fraction corresponding to intact V-ATPase was kept and flash frozen. For negative stain electron microscopy, a single batch of purified V-ATPase was used. For cryoEM, ten batches of purified protein, corresponding to 80 lemons, were thawed and concentrated to 3 μL with a 100 kDa MWCO concentrator prior to freezing specimen grids.

### Mass spectrometry

Samples were prepared for mass spectrometry and mass spectrometry experiments were performed with standard service protocols at the Network Biology Collaborative Centre of the Lunenfeld-Tanenbaum Research Institute. Briefly, samples were reduced with 20 mM dithiothreitol for 10 min at 95°C and alkylated with 40 mM iodoacetamide for 30 min in the dark. Sodium dodecyl sulfate was added to 5 % and phosphoric acid to 1.2 %. S-Trap protein binding buffer (165 μL), consisting of 90 % methanol and 100 mM tetraethylammonium bromide (TEAB) was added of to the acidified lysate (27.5 μL). The resulting mixture was passed through the micro column at 4000 ×g and washed 4 times with the S-Trap protein binding buffer. The sample was digested with 1 μg of trypsin (in 20 μl of 50 mM TEAB) for 1 h at 47°C. Prior to elution, 40 μL of 50 mM TEAB (pH 8) was added to the column and peptides were eluted by centrifugation at 4000 ×g. Remaining peptides were eluted with two 40 μL additions of 2 % formic acid 40 μL) and 50 % acetonitrile with 2 % formic acid. Eluted peptides were dried and stored at -40°C. For data-dependent acquisition, digested peptides were analyzed with a nano-HPLC (High-performance liquid chromatography) coupled to MS. One-tenth of the sample was used for DDA. Coated Nano-spray emitters were generated from fused silica capillary tubing, with 75 μm internal diameter, 365 μm outer diameter, and 5-8 μm tip openings, using a laser puller (Sutter Instrument Co., model P-2000, with parameters set as heat: 280, FIL = 0, VEL = 18, DEL = 2000). Nano-spray emitters were packed with C18 reversed-phase material (Reprosil-Pur 120 C18-AQ, 1.9 μm) resuspended in methanol using a pressure injection cell. The sample in 5 % formic acid was loaded directly at 400 nL min^-1^ for 20 min onto a 75 μm × 15 cm nano-spray emitter. Peptides were eluted from the column with an acetonitrile gradient generated by an Eksigent ekspert™ nanoLC 425 and analyzed on an Orbitrap Fusion™Lumos™Tribrid™. The gradient was delivered at 200 nL min^-1^ from 2.5 % acetonitrile with 0.1 % formic acid to 35 % acetonitrile with 0.1 % formic acid using a linear gradient of 90 min. This elution was followed by an 8 min gradient from 35 % acetonitrile with 0.1 % formic acid to 80 % acetonitrile with 0.1 % formic acid. Next, an 8 min wash with 80 % acetonitrile and 0.1 % formic acid was performed, and equilibration was carried out for another 23 min to 2.5 % acetonitrile with 0.1 % formic acid. The total DDA protocol length was 150 min. The MS1 scan had an accumulation time of 50 ms within a mass range of 400–1500 Da, using orbitrap resolution of 120000, 60% RF lens, AGC target of 125%, and 2400 volts. This process was followed by MS/MS scans with a total cycle time of 3 s. An accumulation time of 50 ms and 33% HCD collision energy was used for each MS/MS scan. Each candidate ion was required to have a charge state from 2-7 and an AGC target of 400 %, isolated using orbitrap resolution of 15,000. Previously analyzed candidate ions were dynamically excluded for 9 s.

### Spectrum matching and de-novo sequencing

Protein identification was carried out with *MaxQuant* version 2.0.3.0 (Cox and Mann, 2008). Spectra were matched against the proteomes of *Citrus clementina* (Cclementina_182_v1.0.protein.fasta, 33929 entries) and *Citrus sinensis* (Csinensis_154_v1.1.protein.fasta, 46147 entries). Proteome sequences were obtained from *Phytozome* (Goodstein et al., 2012). In *MaxQuant*, carbamidomethylation of cysteine was set as a fixed modification; oxidation of methionine as well as phosphorylation of serine, threonine and tyrosine were set as variable modifications. Mass tolerance for the database search was set to 20 ppm on full scans and 0.5 Da for fragment ions. Peptide false discovery rate (FDR) and protein FDR were set to 0.01. Hits to contaminants (e.g. keratins) and reverse hits identified by *MaxQuant* were excluded from further analysis. In addition to matching spectra to existing lemon family proteomes, spectra were sequenced de-novo with the *MaxNovo* algorithm (Gutenbrunner et al., 2021) with default settings. Thus, each matched spectrum to the citrus proteomes also came with a de-novo interpretation that was used for structure interpretation. Subunits of V-ATPases were annotated among the identified proteins as orthologs of respective *Arabidopsis* V-ATPase subunits. The msms.txt output file from *MaxQuant* was then used to filter for V-ATPase subunit peptides. 191 different protein sequences were identified matching to 11 V-ATPase orthologs from *Arabidopsis*.

### CryoEM specimen preparation and data collection

Purified V-ATPase (1.2 μL) was applied to a home-made nanofabricated (Marr et al., 2014) holey gold grid (Meyerson et al., 2014; Russo and Passmore, 2014), that was previously glow-discharged in air for 2 min, in the tweezers of a Leica EM GP2 grid freezing device (>80% RH, 277 K). Excess protein was removed by blotting for 1s before plunge freezing in liquid ethane. The grid was clipped into an auto grid and transferred to a Titan Krios G3 electron microscope (Thermo Fisher Scientific) operating at 300 kV and equipped with a Falcon 4 camera. High-throughput data collection was automated with the *EPU* software package. Movies consisting of 29 exposure fractions were collected at a nominal magnification of 59,000 ×, corresponding to a calibrated pixel size of 1.3475 Å. A total exposure of ∼31 electrons Å^-2^ was used and 12,974 movies were collected.

### Image analysis

Movies were aligned with UCSF MotionCor2 (Zheng et al., 2017) through *Relion* v3 (Scheres, 2012) and processed using *cryoSPARC* v3. CTF parameters were estimated in patches and template selection of particle images was performed. 2D classification with *Relion* v3 was then used to remove undesirable particle images. Multiple rounds of *ab initio* refinement were used to further clean the dataset and a consensus 3D map at 3.8 Å resolution from 189,756 particle images was calculated with non-uniform refinement (Punjani et al., 2020). This consensus set of particle images, with their corresponding orientation parameters, were transferred to *Relion* v3 for particle polishing (Zivanov et al., 2019) and were separated into two distinct rotational states using 3D classification. These particle images (76,072 for State 1 and 104,874 for State 2) were transferred back to *cryoSPARC* v3 for non-uniform refinement, resulting in a State 1 map with a global resolution of 4.0 Å and a State 2 map with a global resolution of 3.9 Å. Focused refinement of the V_1_ and V_O_ regions improved the local resolutions of these areas. A consensus map combining these focused-refined regions was used for subsequent model building. 3D variability analysis in *cryoSPARC* v3 was performed on the particle images that contributed to the highest-resolution maps in each rotary state while masking subunit H with a loose mask. The particle imagess were then subjected to 3D classification using the same mask, revealing the two conformations of H in each rotary state.

### Atomic model building

To begin model building for the high-resolution model (lacking the C-terminal domain of H), homology models were prepared for all subunits from existing structures. For all subunits except AP2, homology models from *PHYRE2* (Kelley et al., 2015) based on yeast V-ATPase models (PDB code 3J9U (Zhao et al., 2015) and 6M0R (Roh et al., 2018)) and models from *RoseTTAfold* (Baek et al., 2021) were compared and the better-fitting model placed in the cryoEM map. In most cases the model were similar or the *RoseTTAfold* model fit better. For AP2, a homology model was built using the rat brain V-ATPase model (PDB code 6VQ7). For both AP1 and AP2 residue identities were removed due to a lack side chain features in the maps.

For the models with subunit H in contact with subunit a, a homology model was prepared for subunit H from yeast subunit H (PDB code 3J9T (Zhao et al., 2015)) with *PHYRE2*. For the models with subunit H in contact with an AB pair, a homology model for subunit H was prepared from the V_1_ΔC crystal structure (PDB code 5D80 (Oot et al., 2016)). The C-terminal domain was then manually rotated and adjusted to fit the density with *UCSF Chimera* (Pettersen et al., 2004).

All models were adjusted manually to fit the experimental maps in *Coot* and refined in multiple rounds with *ISOLDE* (Croll, 2018), *phenix*.*real_space_refine* (Adams et al., 2010), and *phenix*.*rosetta_refine* (DiMaio et al., 2013). Models were validated with *PHENIX* and the PDB online validation tool (wwPDB consortium, 2019).

## Contributions

YZT purified the protein, performed cryoEM experiments, and performed image analysis. KAK performed image analysis and contributed to building of the atomic models, and refined the atomic models. RA built an initial atomic model. WS analyzed mass spectrometry data. PI acquired funding to support RA. KS coordinated experiments and analysis of results. JLR conceived of the study and coordinated and supervised experiments. KAK and JLR wrote the manuscript and prepared the figures with input from all the other authors.

## Acknowledgements

We thank Yazan Abbas for suggestions on image analysis, Stuart Ferguson for discussions about bioenergetics, and Samir Benlekbir for assistance with data collection. YZT was supported by a postdoctoral fellowship from the Canadian Institutes of Health Research, The Agency for Science, Technology and Research Singapore, National University of Singapore (NUS) Presidential Young Professorship, and Ministry of Education Singapore; and JLR was supported by the Canada Research Chairs program. This research was supported by a Natural Sciences and Engineering Research Council Discovery Grant (JLR), and Deutsche Forschungsgemeinschaft grants SFB1101 and TPA02 (KS). Cryo-EM data was collected at the Toronto High-Resolution High-Throughput cryo-EM facility supported by the Canada Foundation for Innovation and Ontario Research Fund. Mass Spectrometry data were collected at the Network Biology Collaborative Centre at the Lunenfeld-Tanenbaum Research Institute, a facility supported by Canada Foundation for Innovation funding, by the Government of Ontario, and by Genome Canada and Ontario Genomics (OGI-139).

## Data Availability

Electron microscopy density maps are available from the electron microscopy data bank with accession codes XXXX-XXXX. Atomic models are available from the protein data bank with access codes XXXX-XXXX.

## Figures

**Supplementary Figure 1.**
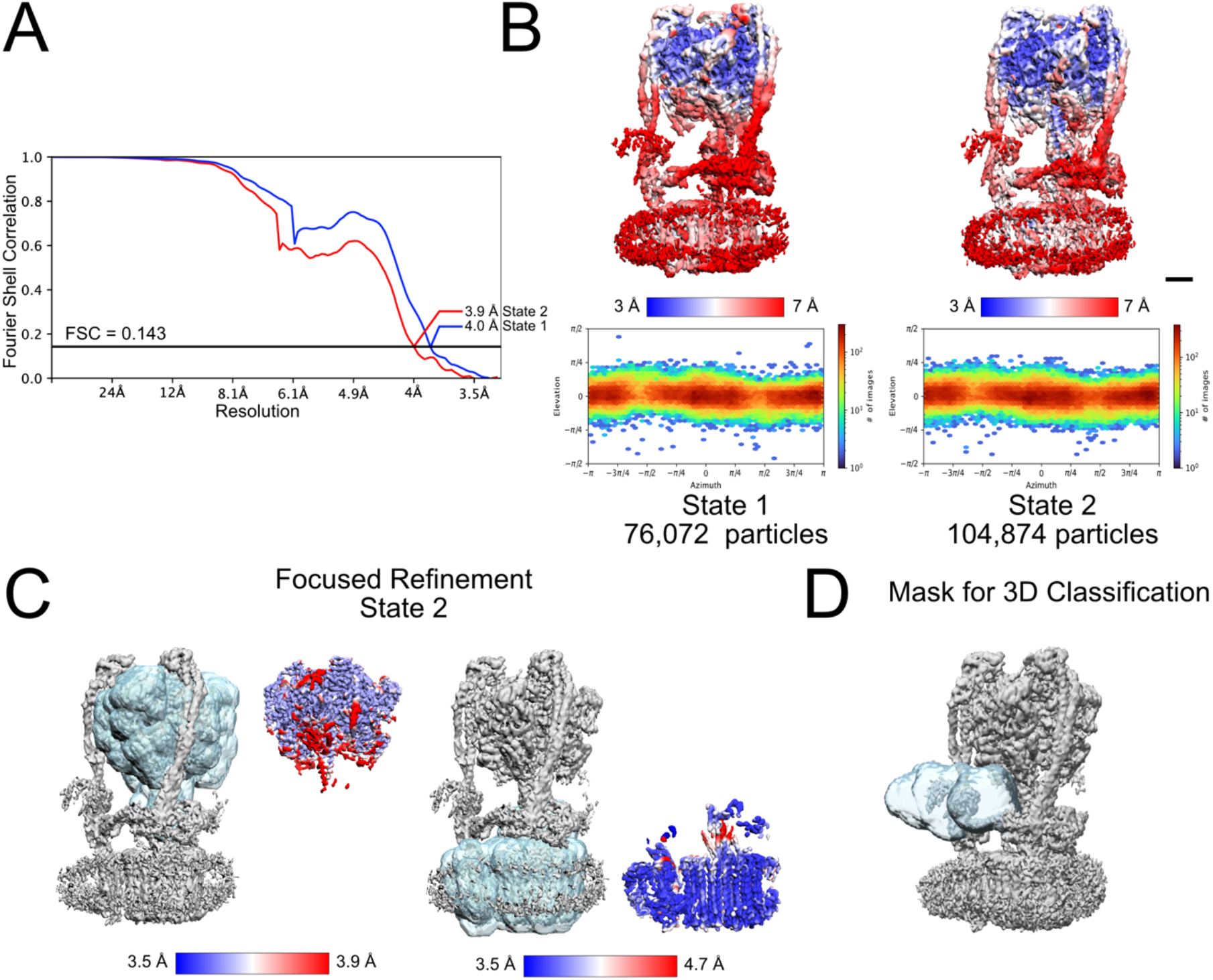
CryoEM map validation. **A**, Fourier Shell correlation following gold standard refinement of the rotational State 1 (*Blue*) and State 2 (*red*) maps, with correction for masking. **B**, Local resolution following global refinement (*top*) and particle orientation distribution (*bottom*) for rotational State 1 (*left*) and rotational State 2 (*right*). **C**, Focused refinement masks (*semi-transparent blue*) and resulting local resolution for the V_1_ region (*left*) and V_O_ region (*right*). **D**, Mask used for local variability analysis and 3D classification of subunit H (*semi-transparent blue*).

**Supplementary Figure 2.**
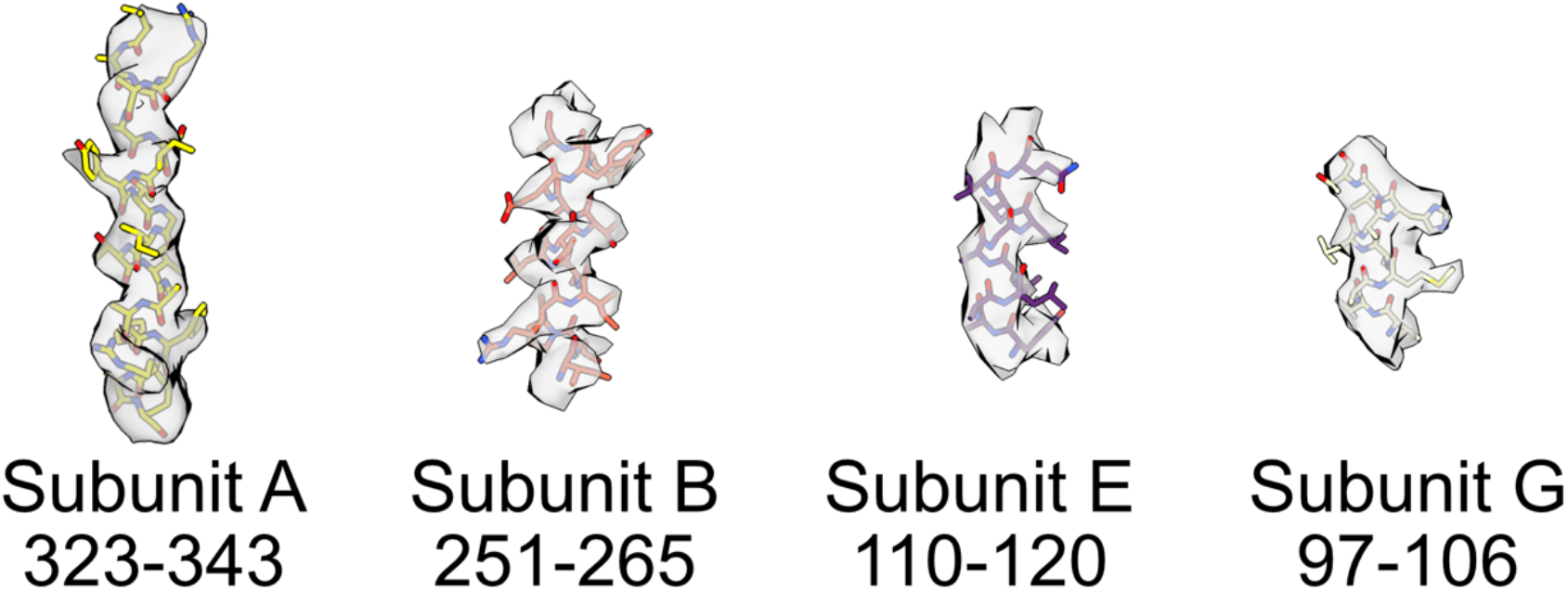
Example model-in-map fit for the highest-resolution regions in the rotational State 2 map.

**Video 1: Model of the citrus V-ATPase in State 2, interpolating between the two conformations of H**.

**Video 2: Model of the citrus V-ATPase, interpolating between State 1 and State 2 with subunit H in contact with subunits A and B**.

**Supplementary Data 1: Raw peptides identified by mass spectrometry**.

